# H-current modulation of cortical Up and Down states

**DOI:** 10.1101/2024.04.05.588281

**Authors:** Leonardo Dalla Porta, Almudena Barbero-Castillo, José Manuel Sanchez-Sanchez, Nathalia Cancino, Maria V. Sanchez-Vives

## Abstract

Understanding the link between cellular processes and brain function remains a key challenge in neuroscience. One crucial aspect is the interplay between specific ion channels and network dynamics. This work reveals a role for h-current, a hyperpolarization-activated cationic current, in shaping cortical slow oscillations. Cortical slow oscillations exhibit rhythmic periods of activity (Up states) alternating with silent periods (Down states). By progressively reducing h-current in both cortical slices and in a computational model, we observed Up states transformed into prolonged plateaus of sustained firing, while Down states were also significantly extended. This transformation led to a five-fold reduction in oscillation frequency. In a biophysical recurrent network model, we identified the cellular mechanisms: an increased input resistance and membrane time constant, increasing neuronal responsiveness to even weak inputs. HCN channels, the molecular basis of h-current, are known neuromodulatory targets, suggesting potential pathways for dynamic control of brain rhythms.

## Introduction

Brain state dynamics are constantly changing in the context of behavior or cognitive demands (e.g., sleep, wakefulness, attention) ^1,2^. Distinct patterns of spatiotemporal dynamics have emerged as brain state correlates, resulting from intrinsic properties of neurons and network connectivity. One of these patterns, cortical slow oscillations provide an illustrative example of this cellular and network interaction.

Slow oscillations are generated by the cerebral cortex in different situations, such as during slow wave sleep (SWS), under deep anesthesia, and even in isolated cortical tissue in vitro, to the extent that they are considered the default activity pattern of the cortical network ^3–5^. Slow oscillations, characterized by interspersed Up and Down states, are thus a hallmark of both intact and isolated cortical networks. Up states, periods of persistent firing, arise from recurrent excitation and network reverberation, while Down states (silent periods) are triggered by the build-up of inhibitory currents during Up states ^6–9^.

The role of slow oscillations during NREM sleep has been associated to different functions such as memory consolidation, synaptic downscaling, homeostasis, or metabolic waste clearance ^10–14^. Slow oscillations are also generated in pathological conditions, such as in perilesional tissue or disorders of consciousness ^4,15–18^. Therefore, understanding the mechanisms of slow oscillations are important not only to explain the healthy brain’s physiological workings and state transitions, but also to deciphering the mechanisms of brain pathologies.

Although considerable progress has been made towards a detailed understanding of the mechanisms controlling slow oscillations, most of these studies focus on synaptic properties and network connectivity ^19–26^. Nonetheless, ion channels also play a fundamental role in shaping brain dynamics, often being targeted by neuromodulators ^1,12,27,28^. One important target of neuromodulators is the hyperpolarization-activated cation current (h-current or *I_h_*) (for a review, see ^29,30^).

H-current is mediated by hyperpolarization-activated cyclic nucleotide–gated (HCN) channels and has a prominent role in rhythmic firing, dendritic excitability, and synaptic integration ^30^. It was first described as a “funny” current (*I_f_*) in cardiac tissue, and later as “queer” current (*I_q_*) in hippocampal cells, due its unique properties: a slow inward current activated by hyperpolarization beyond approximately −50mV to −70mV, carried by Na^+^ and K^+^ ions, that slowly depolarizes the membrane toward its equilibrium potential of ca. −30mV (reviewed in ^29,30^. In the thalamic system, h-current has been well documented as shaping rhythmicity ^31–34^. However, its effect on cortical slow oscillations remains poorly explored.

In this study, we investigate the impact of h-current on spontaneous cortical slow oscillations. Slow oscillations serve as an ideal testbed due to their network-generated nature, integrating both intrinsic neuronal and synaptic properties. Employing a detailed biophysical computational model of the cortical network circuitry allowed us exploring a broader parameter space, in order to identify the mechanisms underlying h-current’s network effects. Our findings reveal the significant modulatory influence of a single ion current on cortical Up and Down state dynamics, strongly influencing properties like the duration of the Up states or persistent firing periods, the Down states and oscillatory frequency. Given that HCN channels, the molecular basis of h-current, are established targets for neuromodulators ^30,35^, our work sheds light on the mechanisms governing the dynamic control of brain rhythms through neuromodulatory pathways.

## Results

To explore the influence of h-current on the dynamics of slow waves in the cerebral cortex, we conducted a series of experiments in cerebral cortex slices *in vitro* and in a computational model. We first recorded spontaneous slow oscillations characterized by alternating periods of neuronal firing or Up states, and periods of silence or Down states. Subsequently, we bath-applied increasing concentrations (10, 50, and 100 µM) of ZD7288 (IC50 = 20 µM;^36^), an h-current antagonist, or an HCN channel blocker ^37–39^. We then analyzed the effects of blocking the h-current on these cortical emergent dynamics generated by the cortical network. Finally, to search more deeply into the mechanisms linking neuronal intrinsic properties to network activity, we investigated the role of h-current in a biophysically realistic Hodgkin-Huxley model of the cerebral cortex network.

Slow oscillations are a prominent activity pattern emerging from the cortical network not only during slow wave sleep, but also under deep anesthesia ^4,40–43^. Interestingly, this activity also emerges in isolated pieces of cerebral cortex such as cortical slices ^6,44–46^, and has been considered the default activity of the cortical network ^47^. This notion is further supported by the finding that slow oscillations can synchronize perilesional areas in the awake brain ^17,18^, potentially reflecting a return to a default state. Figure 1 illustrates spontaneous slow oscillations in a cortical slice, revealed by local field potential and multiunit activity. In this case, the spontaneous rhythmic pattern appears with a frequency of 0.89±0.40 Hz, while Up and Down states are indicated by pink and gray boxes respectively (Fig. 1a). In this same slice, we illustrate the transformation of Up and Down states as a result of progressive h-current block by 10, 50 and 100 µM of the antagonist ZD7288. Up states, which are a period of persistent activity generated by the cortical network recurrency ^8,48,49^ became significantly longer as a result of blocking h-current, going from an average control duration of 0.38±0.18 s to a firing plateau of 2.27±1.21s in 100 µM ZD7288. The periods of silence of each oscillatory cycle or Down states also were remarkably elongated, from 1.96±2.20 s at baseline to 10.97±6.98 s in 100 µM ZD7288. This profound transformation of the emergent activity resulted in a decrease in oscillatory frequency, as a consequence of longer Up and Down states, which can be observed in the autocorrelogram (Fig. 1c). The baseline frequency of 0.89±0.40 Hz decreased to 0.11±0.10 Hz in 100 µM ZD7288.

**Fig. 1.**
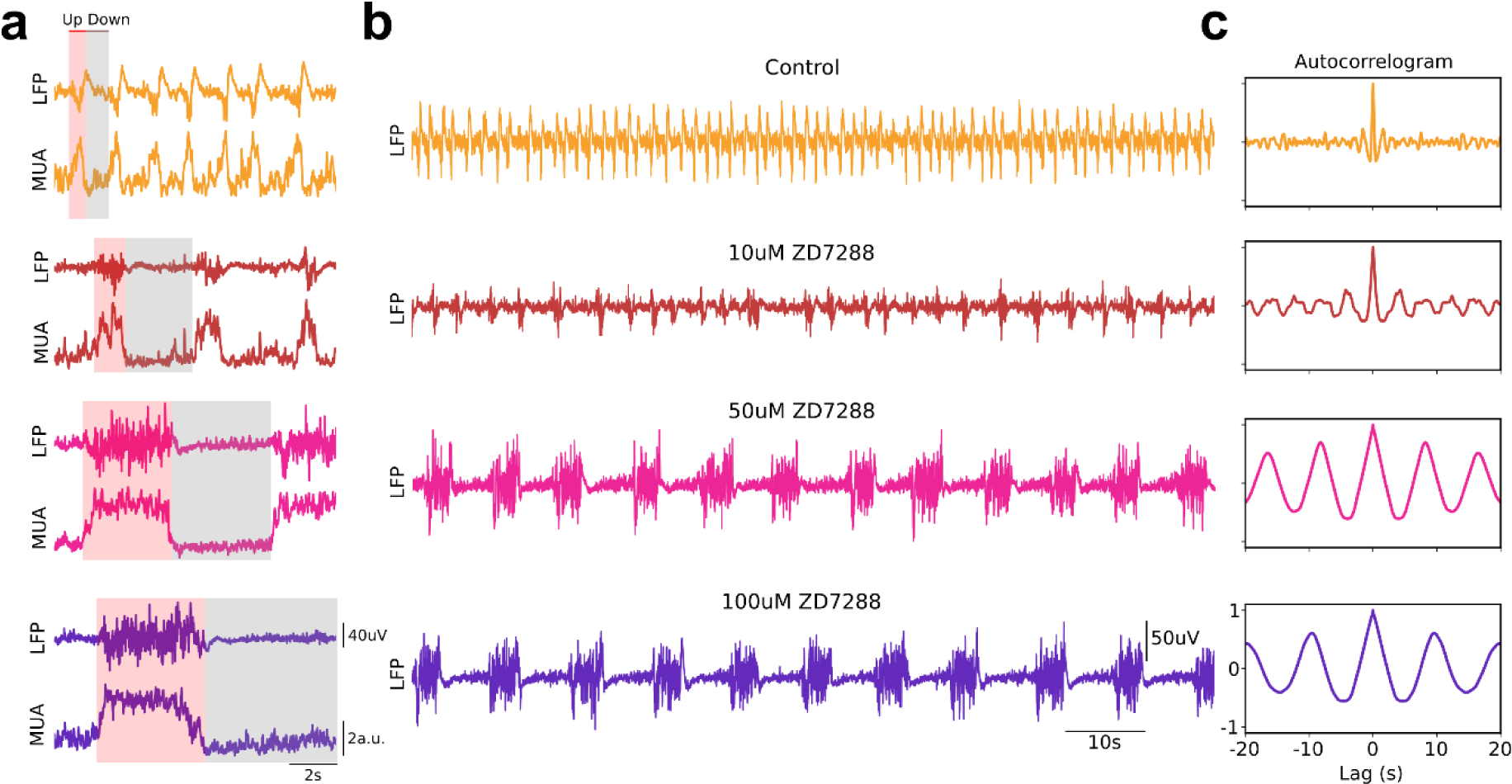
h-current blocker ZD7288 effects on spontaneous slow oscillations. **a.** Spontaneous Up and Down states, which were detected from the relative firing rate (MUA). **b.** A hundred seconds of raw local field potential (LFP) illustrating network activity during SO in cortical slices and after bath application of ZD7288 in three different concentrations: 10, 50 and 100 µM, respectively. **c.** Autocorrelograms of the activity displayed in b for a 40 s window. The resulting oscillatory frequencies were 0.89±0.40 Hz, 0.36±0.25 Hz, 0.15±0.22 Hz and 0.11±0.10 Hz for control and 10, 50 and 100 µM ZD7288, respectively.

The temporal profile of the elongation of Up and Down states is demonstrated in the average firing rate across detected Up states and centered around Down to Up transitions (Fig. 2a). In it, the prolongation of the Up states goes from a period of firing of under 1 s in the control condition, to a prolonged plateau of persistent firing lasting for over 3 s when the h-current is blocked.

**Fig. 2.**
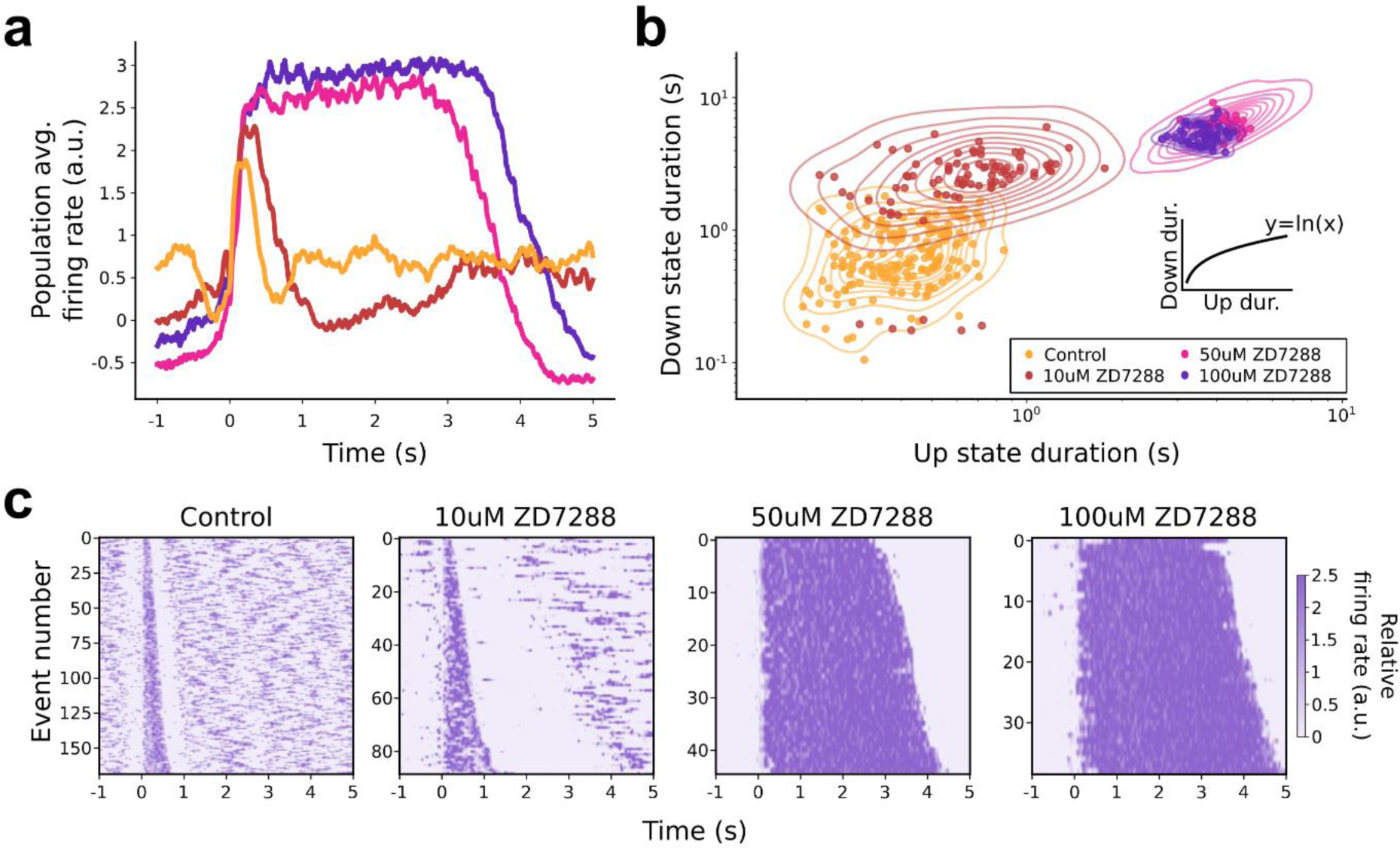
Up and Down states modulation by h-current antagonist. **a.** Population average relative firing rate (MUA) across detected Up states and centered around Down to Up transitions. **b.** Scatter plot of Up and Down state durations for control slow oscillations and after bath-application of different concentrations of ZD7288 (see inset), a h-current antagonist. Irregular ellipses represent the bivariate (2D) kernel density estimated (KDE). Up states were detected in a 200s-long time series and the number of Up states detected were 169, 89, 45 and 39 for control and 10, 50 and 100 µM ZD7288, respectively. **c.** Raster plot of relative firing rate (MUA) for all Up and Down states detected in each condition (200s-long time series). The data corresponds for an illustrative single case, same as in Fig. 1.

A more in-depth analysis of the elongation of Up and Down states resulting from the h-current block is presented in Fig. 2b. A scatter plot represents the individual oscillatory cycles according to their Up and Down states duration in the same slice shown in Fig. 1 at baseline and in 10, 50, and 100 µM of ZD7288. Up and Down states are thus dynamically related, such that the Down states are highly dependent on the firing rate frequency and duration during Up states. In other words, Down states have an activity-dependent component which can be mediated to a large extent by hyperpolarizing currents, probably potassium currents. This dynamical link has been demonstrated both experimentally ^24,26,28^ and computationally ^8,50^. This relationship is also observed here, not only in the scatter plot but also in the raster plots (Fig. 2c), where we display individual events ordered by the duration of the Up states during a window of 200 s for control and 10, 50 and 100 µM ZD7288, respectively. We quantified this relationship by fitting the points on the scatter plot, which yielded the following functional relationship:

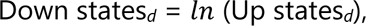

where Down states*_d_* and Up states*_d_* are the duration of Down and Up states, respectively. Therefore, the duration of Down states increases with the duration of Up states, albeit at a slower rate, an observation that holds for the studied population.

As shown for the example in Figs 1 and 2, blocking the h-current with ZD7288 resulted in prominent changes in the spontaneous emergent activity from the cortical network in the population average (Table 1). The application of 10 µM ZD7288 resulted in a 197% elongation of the Up state duration (to 0.75±0.45 s) and a 246% elongation of the Down state duration (to 4.83±4.59 s; Fig. 3a,b). Higher doses of ZD7288 (100 µM) resulted in a further elongation of the Up state duration to 2.27±1.21s and to a Down state duration of 10.97±6.98 s. As a result, the oscillatory frequency decreased from 0.53±0.28 Hz to 0.11±0.16 Hz with the increasing concentration of h-current blocker resulting in an 83% decrease in network frequency for 100µM of ZD7288 (0.11±0.16 Hz; Fig. 3c).

**Fig. 3.**
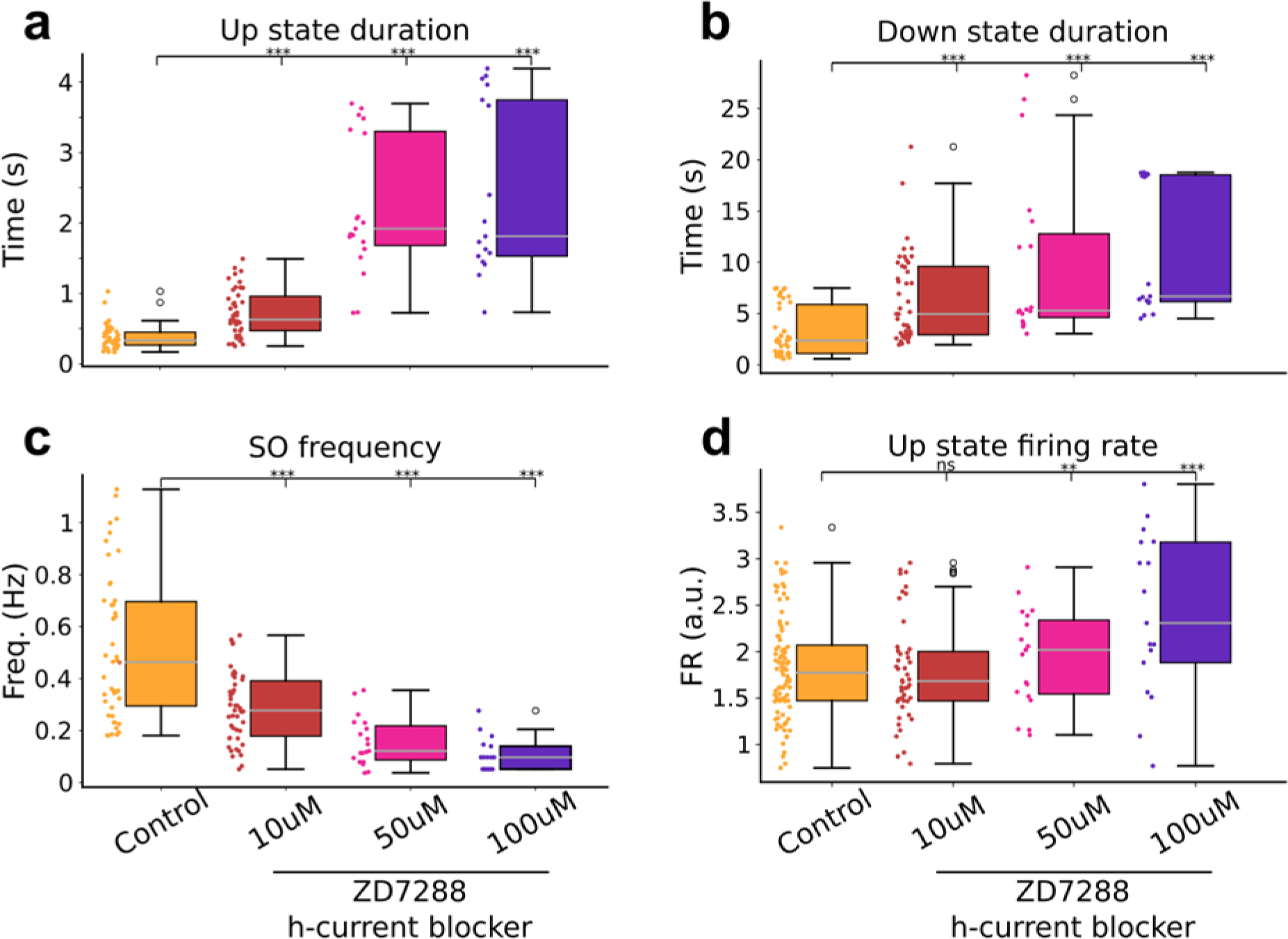
Up and Down states variation with h-current block. **a.** Duration of Up states, **b.** duration of Down states, **c.** slow oscillations frequency, and **d.** firing rate during Up states, in control and three levels of h-current block induced by 10, 50 and 100 µM ZD7288. ** *p*≤0.01; *** *p*≤0.001; ns = not significant; (two-sided Wilcoxon signed-rank test).

**Table 1.** Relative changes of Up and Down state properties during blockage of h-current. *P*-value of a two-sided Wilcoxon signed rank test (\*\**p*<0.01, \*\*\**p*<0.001; ns=not significant)

| **Parameter** | **Control**<br>(*n*=13) | **10uM ZD7288**<br>(*n*=13) | **50uM ZD7288** (*n*=8) | **100uM ZD7288**<br>(*n*=6) |
| --- | --- | --- | --- | --- |
| Frequency (Hz) | 0.53±0.28 | 0.33±0.39(***) | 0.20±0.26(***) | 0.11±0.16(***) |
| Up state duration (s) | 0.38±0.18 | 0.75±0.45(***) | 2.08±1.26(***) | 2.27±1.21(***) |
| Down state duration (s) | 1.96±2.20 | 4.83±4.59(***) | 6.35±5.88(***) | 10.97±6.98(***) |
| Up state relative firing rate<br>(a.u.) | 1.93±0.46 | 1.77±0.50 (ns) | 2.04±0.42 (**) | 2.55±0.76 (***) |

With respect to the population firing rate during Up states, it did not increase significantly in 10µM ZD7288, but increased by 32% (from 1.93±0.46 a.u. to 2.55±0.76 a.u.) in 100µM ZD7288 (Fig. 3d). Such an increase in firing rate is intriguing, since the blockade of h-current results in a hyperpolarization of the neuronal membrane potential. This is the case given that an open h-current depolarizes towards its reversal potential (−30 mV; ^29^). How the hyperpolarization of the membrane potential can lead to a population increase in the firing rate is something that we explore in the spiking network model in the next section.

Our study so far has provided experimental evidence that modulating a particular ionic channel, the HCN channel, which is an inherent cellular characteristic, significantly affects the emergent patterns observed in neuronal networks. This effect is likely amplified by the recurrent connectivity characteristic of the cerebral cortex ^51–53^. To elucidate the mechanisms underlying this transition from cellular to network-level effects, we employed a biophysical detail model of the cerebral cortex with recurrent connections, allowing for a detailed dissection of the contributing factors.

### Biophysical network model and h-current

To investigate the underlying mechanisms through which the h-current modulates cortical slow oscillations, we implemented a biophysically detailed computational model of the cortical network ^8,28^. The model comprises pyramidal (*n*=1024) and inhibitory (*n*=256) conductance-based neurons evenly spaced along a line, connected with a probability that decays with the distance between neurons (Fig. 4a). Biologically plausible synaptic dynamics were modeled as excitatory AMPA and NMDA, and inhibitory GABA_A_. The dynamic of the h-current was implemented according to the description in ^54^. H-current was modeled in only 30% of randomly selected pyramidal neurons and was located at the dendritic compartment ^55,56^. Our implementation allowed us to have a heterogeneous network with cell-to-cell variability ^57,58^.

**Fig. 4.**
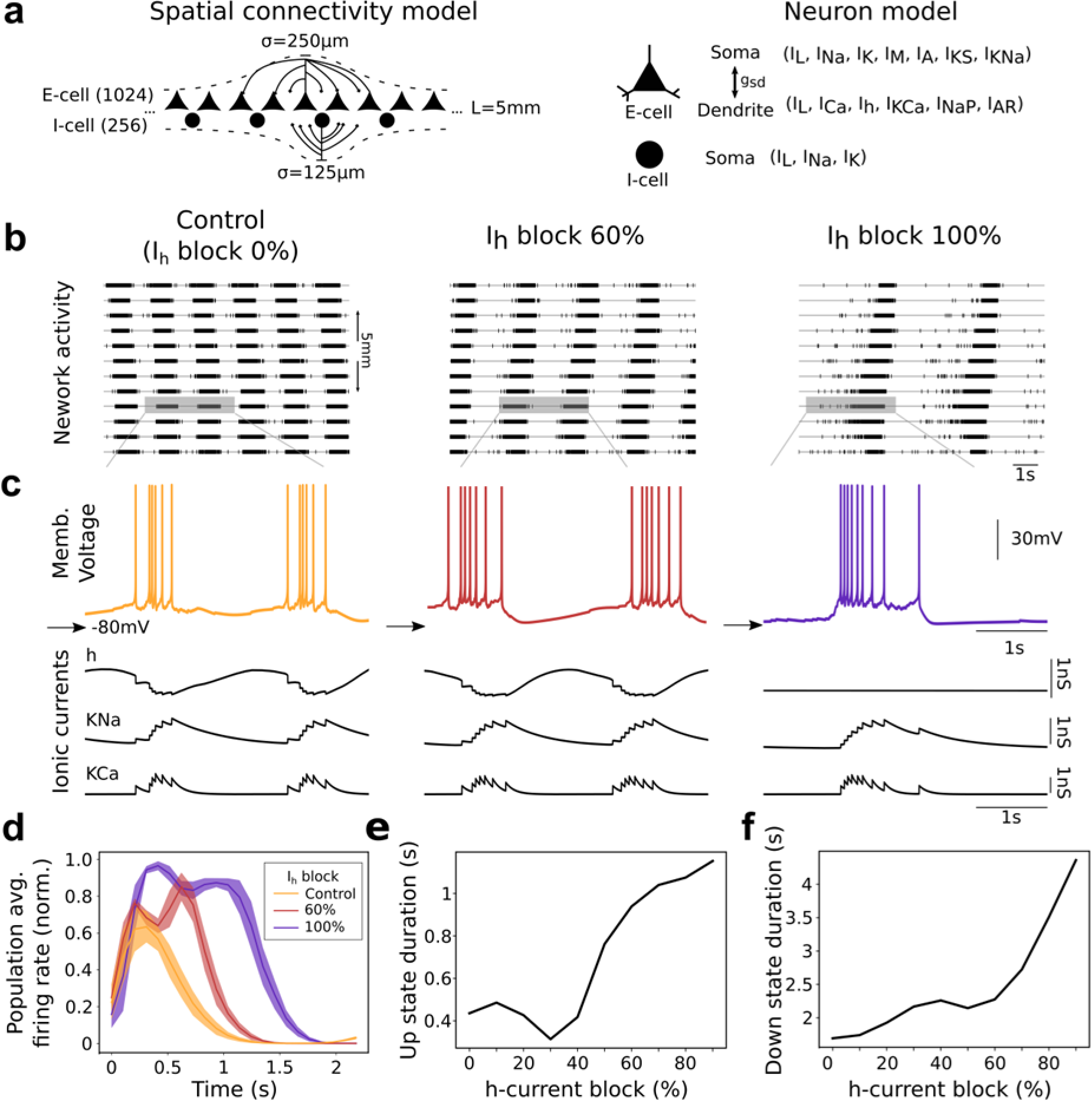
h-current on the cortical network model. **a.** Cellular ionic currents and connectivity in the computational model (see Methods). **b.** Spontaneous network activity visualized as multiunit activity (MUA; 100 neighboring cells per site) during control slow oscillations (left) and with 60% and 0% of h-current block, respectively. **c.** Representative membrane potential of a pyramidal neuron, for the same conditions described in b, and the time evolution of the h-current as well as the sodium-(Na) and calcium-(Ca) dependent potassium currents. **d**. Population average neuronal firing rate (normalized by the maximum) across detected Up states and centered around Down to Up transitions. **e**. and **f**., Up and Down state durations as a function of h-current block, respectively.

In our computational model of the local cortical circuit, we were able to reproduce the slow oscillation network dynamics in the form of Up (active) and Down (silent) states and reproduce the experimental observations (Fig. 4). In the control condition, the slow oscillations occurred with a frequency of 0.62 Hz, with an Up-state duration of 0.44±0.04 s and Down-state duration of 1.68±0.05 s, and with an average neuronal firing rate of 9 Hz (Fig. 4b). Looking into the intracellular dynamics during control conditions (Fig. 4c), we observed that during the Up states, and due to neuronal firing, adaptation builds up (*I_KNa_* and *I_KCa_* in the model) while h-current reaches its minimum due to the membrane potential depolarization (Fig. 4c, left).

Conversely, after an Up state, the membrane is hyperpolarized due to adaptation and the h-current starts to increase while adaptation currents decrease over time. While *I_KNa_* and *I_Kca_* build up the Down state, their progressive reduction goes in parallel with the opening of h-current, and all these events contribute to the triggering of the next Up state. With the block of h-current at 60% and 100% (Fig. 4c middle and right columns), Up and Down states became longer and the number of spikes per Up state increased.

At the population level, we evaluated the impact of h-current expression on the Up and Down states by parametrically varying the expression of h-current. The population average firing rate revealed not only an elongation of Up state duration but also an increment in the neuronal firing rate after blockage of h-current (Fig. 4d). Also, we observed an almost linear relationship between h-current block and the duration of Up and Down states, such that the lower the expression of h-current, the longer the duration of Up and Down states (Fig. 4e, f). For example, during control conditions the duration of Up and Down states were 0.44±0.04 s and 1.68±0.05 s, respectively. While, for a total blockage of h-current, there was a 260% elongation of Up state duration (1.15±0.05 s) (Fig. 4e) and a 258% elongation of Down state duration (4.35±0.07 s) (Fig. 4f).

We next evaluated the impact of h-current at the neuronal level, to understand how it induces changes at the population level. The *f*–*I* relationship for different levels of h-current revealed a modulation of neural gain (Fig. 5a, left). For instance, with a depolarizing injected current of 0.4 nA, the neuron with 100% expression of h-current (control) did not show any spikes. Conversely, with the same injected current, a neuron with 60% block of h-current fired at a frequency of 6 Hz (Fig. 5a), illustrating the impact that h-current has on the neuronal sensitivity to inputs. This increased responsiveness in neurons when h-current is blocked can be explained by the increase in input resistance (Fig. 5a, right side), and it leads to a progressive firing rate across the range of injected current explored (Fig. 5b).

**Fig. 5.**
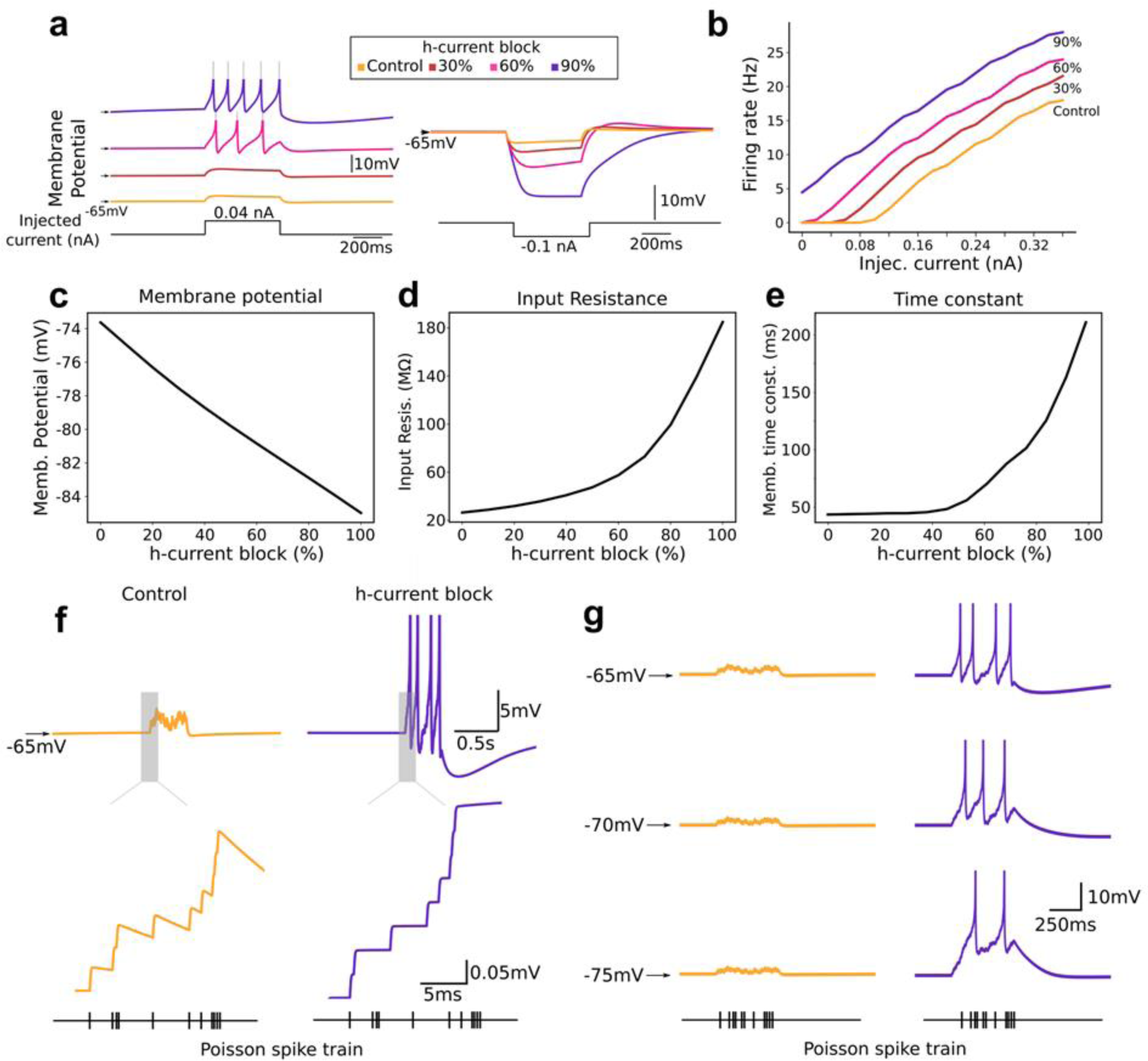
Effects of h-current at the cellular level in the computational model. **a.** Left: neuronal response to an injected depolarizing current of 0.04 nA during 0.5 s for different levels of h-current block. Right: neuronal response to a hyperpolarizing current of −0.1 nA during 0.5 s. Color code represents different h-current block. **b.** Neuronal firing rate as a function of injected depolarizing current (f–I) for different h-current block. Firing rate was defined as the average neuronal rate during a pulse of 0.5 s. **c.** Resting membrane potential, input resistance (**d**), and time constant (**e**) as a function of h-current block. **f**. Single neuron response to a depolarizing Poisson spike train (0.5 Hz rate during 0.5 s) for control (yellow) and full h-current block (purple), respectively. **g**. Same as in (**f**) for different resting membrane potential.

The parametric variation of h-current activation levels was investigated in relation to the neuronal intrinsic properties: membrane potential (Fig. 5c), input resistance (Fig. 5d), and time constant (Fig. 5e). While input resistance and time constant display an exponential growth with h-current blockade, the membrane potential linearly hyperpolarizes. As shown above, the increase in input resistance is critical for the increased responsiveness of neurons to the same inputs. The model reveals that this increase in the response to inputs persists despite the hyperpolarization of the membrane potential. To investigate the cellular to network mechanisms, we simulated the injection of trains of presynaptic action potentials with a Poisson distribution (Fig. 5f). The inputs evoked excitatory postsynaptic potentials, which had larger amplitudes under h-current block. Furthermore, the increased time constant of the membrane resulted in a more efficient summation of synaptic potential, both resulting in a train of postsynaptic action potentials, while with active h-current the response remained below threshold. We then investigated for the corresponding hyperpolarization (5–10 mV) (Fig. 5g) and found that the effect on input resistance and time constant of the membrane could overcome the more negative membrane potential, still presenting a suprathreshold response with a similar number of action potentials. This explains the effect that we have described in the model at the network level, where the recurrency of the cortical network amplifies these effects.

Together, our results suggest that h-current is highly implicated in the control of Up and Down dynamics through modulation of neuronal properties that are transferred to the network by virtue of network recurrency (Fig. 6).

**Fig. 6.**
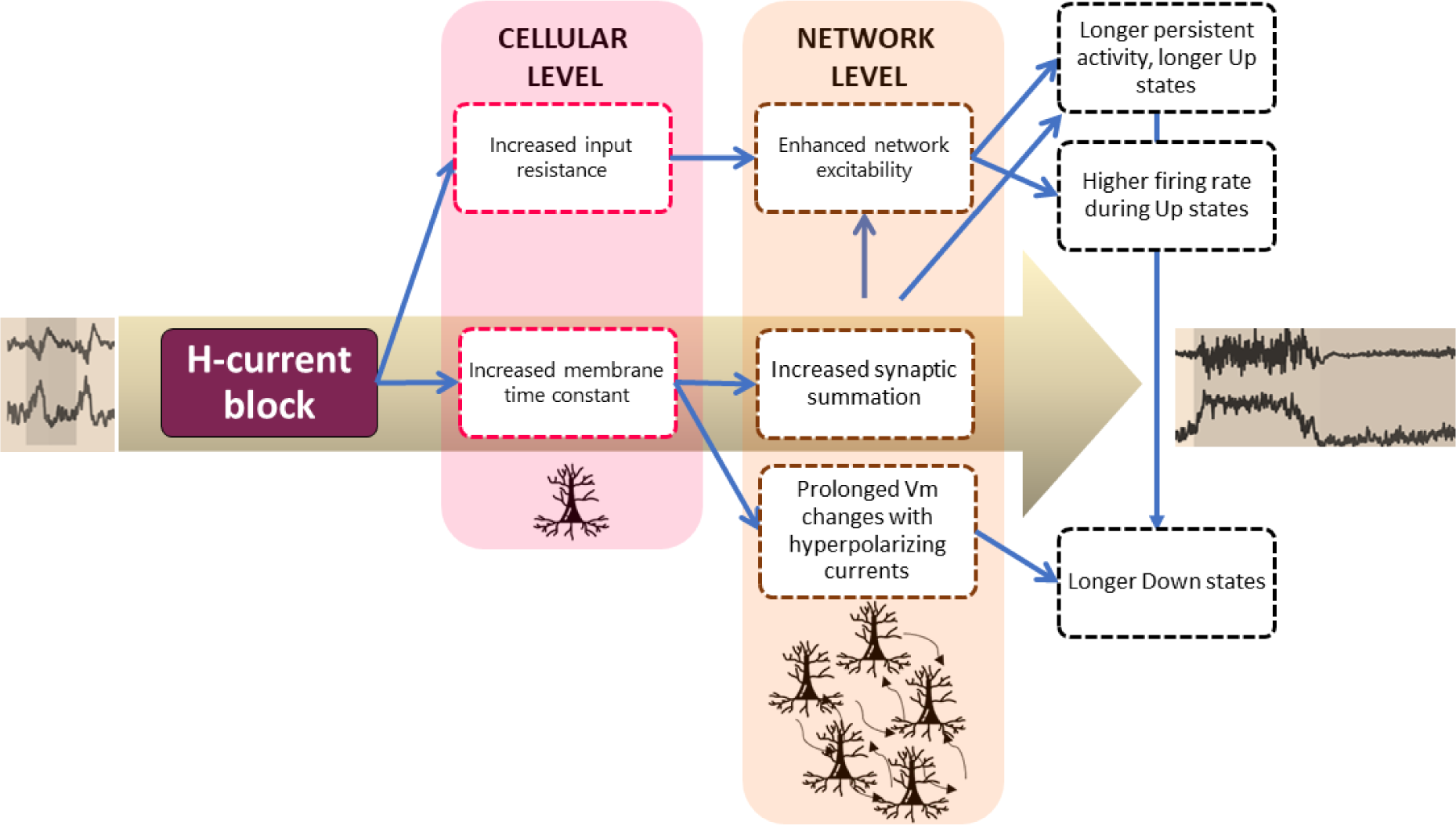
h-current role, from cellular to network level: schematic synopsis. Following h-current block, the increased input resistance and time constant at the cellular level, amplify the amplitude and duration of the synaptic inputs received by the neurons. This is further amplified at the network level, due to the recurrent connections between cortical neurons, resulting in longer Up states with more persistent activity, higher firing rates and larger firing rates, thus inducing longer Down states.

## Discussion

Cortical slow oscillations have been associated with both physiological and pathological processes, yet the underlying mechanisms remain unclear. Here, we investigated the correlates of h-current (*I_h_*) during slow oscillations, both *in vitro* and *in silico*, and showed that h-current had a pronounced effect on cortical Up and Down dynamics. We demonstrated that h-current not only modulates the population neuronal firing rate but also controls the duration of both Up and Down states. Further, in a detailed biophysical recurrent model of the cortical network, we demonstrated that the observed population effects largely depend on the impact of h-current at the cellular level. Intrinsic neuronal properties, such as input resistance and membrane time constant, were modulated by h-current, facilitating more efficient summation of synaptic potentials, thereby mechanistically explaining the experimental results. Therefore, the present findings link the role of specific ion channels and collective network behavior during cortical slow oscillations (for an overview, see Fig. 6).

Slow oscillations are a key feature of the cerebral cortex and are characterized by depolarized periods of sustained firing (Up states) followed by hyperpolarized periods of silence (Down states or off-periods), oscillating at a frequency of approximately 1 Hz ^40,59^. Slow oscillations emerge spontaneously during both physiological and pathological conditions ^5^. During NREM sleep, slow oscillations emerge locally, most prominently in cortical layer 5, and propagate throughout the cortex as a travelling wave ^6,60^. Its functional benefits have been linked to brain clearance, cellular maintenance, and memory consolidation ^12,14,61,62^. On the other hand, slow oscillations also manifest in clinical conditions, emerging as a result of pathological states ^4,15^, or around brain lesions ^17,18^. Therefore, understanding the underlying mechanisms behind slow oscillations is not only crucial for comprehending the physiological functioning of the brain and its transitions, but is also vital for understanding pathological processes within the brain.

Here we investigated the cellular substrates of cortical slow oscillations, revealing h-current as a key modulator of the collective neuronal behavior (Fig. 1). H-current was first described as a pacemaker current, because of its involvement in the generation and control of spontaneous activity in cardiac cells. Today, it’s known as a voltage-dependent current highly selective for Na^+^ and K^+^, mediated by the HCN channels that are located throughout the brain (for a review see ^29,30^).

In the thalamocortical system, h-current has been described as being involved in the generation of slow periodic rhythms such as spindle waves ^32,33,38,63^. At the cellular level, h-current has been shown to modulate neuronal excitability, resting membrane potential, neuronal impedance, response to hyperpolarization, resonance properties, sequential neuronal patterns, as well as contributing to oscillatory frequency ^33,39,58,64–68^.

However, despite its prominent role in slow frequency oscillations, h-current has been mainly explored in the thalamic system. Its effect on cortical slow oscillations remained limited, with the exception of a recent study by Shu et al. ^69^, where they described the role of h-current in controlling recurrent persistent activity (Up states), a factor attributed to the increase in input resistance upon blockade of h-current.

Here, we found that h-current has a remarkable role in the mechanisms controlling not only the periods of persistent activity (Up state) and neuronal firing, but also the periods of neuronal silence (Down state). Blocking h-current with an h-current antagonist (ZD7288) at various doses resulted in prominent modulation of both Up and Down state durations (Figs 1–3), revealing a dynamic link among them, such that the Down states were highly dependent on the firing rate frequency and duration during Up states (Fig. 2). In other words, the Down state duration increased with the duration of Up states, albeit at a slower rate. This can be partially explained because Down states have an activity-dependent component which can be mediated largely by hyperpolarizing currents, probably potassium currents ^6,9^. Indeed, previous experimental and computational studies have revealed a dynamic link among Up state and Down state durations ^8,24,26,28^. Despite the strong effect of h-current blocking, we cannot rule out the possibility that other ionic currents may also be involved in this process, since ZD7288 has been reported to affect both Na^+^ channels and T-type Ca^2+^ channels ^70,71^.

To bridge the link between intrinsic neuronal properties (h-current) and population dynamics we extensively explored a biophysically detailed model of cortical slow oscillations ^8^. Apart from reproducing our experimental observations at the population level (i.e., elongation of Up and Down states), we revealed the h-current mechanisms at the neuronal level (Fig. 4). In the model, h-current effectively shaped neuronal intrinsic properties, such as membrane potential, input resistance, and time constant (Fig. 5). The increment in the input resistance, generated by the h-current block, was crucial for an enhancement of the neuronal responsiveness to the same inputs, which persisted even for the hyperpolarization of the membrane potential (Fig. 5). Another crucial fact to explain the effects observed in population dynamics was the modulation of membrane time constant. As h-current blockade increased the membrane time constant, it allowed for a more efficient summation of synaptic potential, which in turn can explain the elongation in the periods of persistence neuronal firing, observed experimentally (Fig. 5).

Together, our experimental and computational findings highlight the profound influence of h-current on cortical network dynamics, as well as the broader interplay between specific ion channels and collective network behavior (Fig. 6). Furthermore, considering that HCN channels, which constitute the molecular basis of h-current, are known targets of neuromodulators, our work provides the mechanisms for the dynamic control of brain rhythms through neuromodulatory pathways ^30,35^.

## Methods

### Ethics statement

Ferrets were treated in accordance with the European Union guidelines on the protection of vertebrates used for experimentation (Directive 2010/63/EU of the European Parliament and of the council of 22 September 2010). All experiments were approved by the ethics committee of the University of Barcelona. Several elements of the methods described here are taken from and described in our previous work ^46^.

### Slice preparation

Ferrets (4–10-months-old; either sex) were deeply anesthetized with isoflurane and sodium pentobarbital (40 mg/kg) before decapitation. The brain was quickly removed and placed in an ice-cold sucrose solution containing (in mM): 213 sucrose, 2.5 KCl, 1 NaH_2_PO_4_, 26 NaHCO_3_, 1 CaCl_2_, 3 MgSO_4_, and 10 glucose. Acute coronal slices (400-µm-thick) of the occipital cortex, including visual cortical areas 17, 18 and 19, from both hemispheres, were extracted.

Slices were placed in an interface-style recording chamber (Fine Science Tools, Foster City, CA) and superfused with an equal mixture of the above-mentioned sucrose solution together with an artificial cerebrospinal fluid (ACSF) as described in ^6^. The ACSF contained (in mM): 126 NaCl, 2.5 KCl, 1 NaH_2_PO_4_, 26 NaHCO_3_, 2 CaCl_2_, 2 MgSO_4_, and 10 glucose. Next, slices were bathed with ACSF for 1–2 h to allow recovery. To obtain spontaneous SO, slices were superfused for at least 30 min before experiments with ACSF containing (in mM): 126 NaCl, 4 KCl, 1 NaH_2_PO_4_, 26 NaHCO_3_, 1 CaCl_2_, 1 MgSO_4_ and 10 glucose. All solutions were saturated with carbogen (95% O_2_ / 5% CO_2_) to a final pH of 7.4 at 34°C.

### Electrophysiological recordings

We recorded the extracellular local field potential (LFP) using a 32-channel multi-electrode array (MEA). The signal was acquired, amplified, and digitized with an ME2100-System and Multi-Channel Experimenter software (Multichannel Systems MCS GmbH-Harvard Bioscience Inc, Reutlingen, Germany), at a sampling frequency of 10 kHz.

### Pharmacological agents

ZD7288 (10, 50, and 100µM), an h-current blocker, was obtained from Tocris Bioscience (Bristol, UK).

### Computational model

For a quantitative investigation and analysis of the effects of h-current in the modulation of SO, we simulated the model of isolated cortical network proposed by Compte et al. ^8^, with the addition of h-current ^28^. The model consists of 1024 pyramidal neurons and 256 inhibitory neurons interconnected through biologically plausible synaptic dynamics. The neurons are equidistantly distributed on a line and sparsely connected to each other, as in the control network described in ^8,28^. Each neuron makes a 20±5 (SD) connection to its postsynaptic targets (autapses are not allowed). The network is assumed to be 5 mm long and the probability of connections between neuron pairs is determined by a Gaussian distribution centered at zero with a prescribed standard deviation. For pyramidal and inhibitory neurons, the standard deviation was set at 250 mm and 125 mm, respectively.

Pyramidal neurons have a somatic and a dendritic compartment. The dynamical equations are:

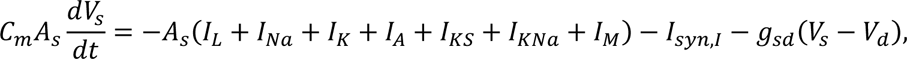

and

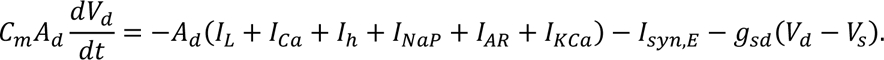

*V*_*s*_(*V*_*d*_) and *A*_*s*_ = 0.015 mm^2^(*A*_*d*_ = 0.035 mm^2^) represent the soma (dendrite) membrane voltage and membrane area, respectively. *C*_*m*_ = 1µ*F*/cm^2^ is the specific membrane capacitance and *g*_*sd*_ = 1.75 ± 0.1µS is the electrical coupling conductance between soma and dendrite. *I*_*syn,E*_ (*I*_*syn,I*_) are the excitatory (inhibitory) synaptic currents. All excitatory synapses target the dendritic compartment, and all inhibitory synapses are localized on the somatic compartment of postsynaptic pyramidal neurons.

The somatic compartment includes the following ionic currents (*I*) and respective maximal conductance (*g*): leakage current (*I*_L_, *g*_L_ = 0.0667 ± 0.0067 mS/cm^2^), sodium current (*I*_Na_, *g*_Na_ = 50 mS/cm^2^), potassium current (*I*_*K*_, *g*_*K*_ = 10.5 mS/cm^2^), A-type K^+^ current (*I*_*A*_, *g*_*A*_ = 0.9 mS/cm^2^), non-inactivating slow K^+^ current (*I*_*Ks*_, *g*_*Ks*_ = 0.403 mS/cm^2^), Na^+^-dependent K^+^ current (*I*_*K*Na_, *g*_*K*Na_ = 0.744 mS/cm^2^), and the non-inactivating and non-rectifying K^+^ current (*I*_*L*_, *g*_*L*_ = 0.083 mS/cm^2^). The dendrite includes leakage current (*I*_L_, *g*_L_ = 0.0667 ± 0.0067 mS/ cm^2^), high-threshold Ca^2+^ (*I*_*C*a_, *g*_*C*a_ = 0.43 mS/cm^2^), non-inactivating hyperpolarization-activated current (*I*_H_, *g*_H_ = 0.0115 mS/cm^2^), persistent Na^+^ channel (*I*_NaP_, *g*_NaP_ = 0.0686 mS/ cm^2^), anomalous rectifier K^+^ channel (*I*_*A*R_, *g*_*A*R_ = 0.0257 mS/cm^2^), and the Ca^2+^-dependent K^+^ current (*I*_*KC*a_, *g*_*KC*a_ = 0.57 mS/cm^2^). The inhibitory neurons, consisting of only a somatic compartment, are modeled as:

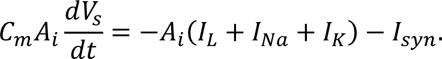

*A*_i_ = 0.02mm^2^ is the total membrane area and *I*_*syn*_is the synaptic current from both the inhibitory and excitatory neurons. The maximal conductances are: *g*_L_ = 0.1025 ± 0.0025 mS/ cm^2^, *g*_Na_ = 35 mS/cm^2^, and *g*_*K*_ = 9 mS/cm^2^. All the details of the implementation of these currents are described in ^8^ except for *I*_ℎ_ and *I*_*L*_, which are described in ^28^. Briefly, the M-current was implemented as described in ^72^: *I*_*L*_ = *g*_*L*_*m*(*V* − *V*_*K*_). The activation variable is controlled by 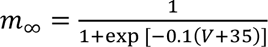 and 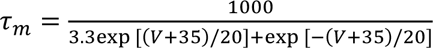. For h-current we implemented the model described by Hill and Tononi ^54^: *I*_ℎ_ = *g*_ℎ_*m*(*V* + 45). The activation variable is controlled by 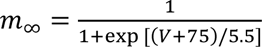 and 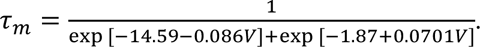. Only 30% of randomly selected pyramidal neurons were endowed with h-current. To simulate the experimental effects of h-current blockage, we progressively reduced the *I*_ℎ_ maximal conductance (*g*_ℎ_) from 100% (control condition) to 0% (which will be referred to as simple expression). The synaptic currents were modeled as described in ^8^ with the following adjustments of AMPA, NMDA and GABA-A maximal conductance: 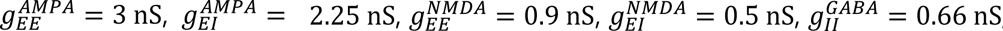, and 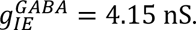.

### Network dynamics analysis

For every condition tested, we analyzed at least 200 s of spontaneous activity. We first estimated multi-unit activity (MUA) from local field potential (LFP) recordings and detected Up and Down states as previously described ^73–75^. The MUA signal was calculated as the average power of the normalized spectra at high-frequency band (200–1500 Hz), since power variations in the Fourier components at high frequencies of LFP provide a reliable estimate of the population firing rate ^76^. The MUA signal was then logarithmically scaled to balance large fluctuations of nearby spikes (log(MUA)). We detected Up and Down states setting duration and amplitude thresholds in the log(MUA) signal. In this way, we could compute different parameters that characterize SO, such as oscillatory frequency or Up and Down state durations. For each Up and Down cycle, the duration for both the Up and Down states were computed. Once extracted, they were scattered one against the other to construct the 2D space of points coloring by condition. Kernel density estimation (KDE) was used to obtain and construct a univariate (1D histogram estimate) and bivariate (2D histogram estimate) plots of the Up and Down state durations ^77^. For the simulated data, the Up states were detected thresholding the Gaussian smoothed network activity (defined as the sum of spikes within a 4 ms bin). In Fig. 4a and b, neuronal membrane potential was held at −65mV to explore the effects of h-current blockage. Input resistance was computed following the Ohm’s law: 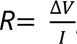, where *V* is the membrane voltage and *I* the applied current (I = −0.1nA for 500 ms).

## Acknowledgements

We thank Tony Donegan for editing.

## Data availability

The code in support of this publication is publicly available at https://github.com/ldallap/M-Current-modulation-of-cortical-slow-oscillations.

